# High Enhancer Activity is an Epigenetic Feature of HPV Negative Atypical Head and Neck Squamous Cell Carcinoma

**DOI:** 10.1101/2021.09.21.461310

**Authors:** S. Carson Callahan, Veena Kochat, Zhiyi Liu, Ayush T Raman, Jonathan Schulz, Christopher Terranova, Margarita Divenko, Archit Ghosh, Ming Tang, Curtis Pickering, Jeffrey N. Myers, Kunal Rai

**Author notes:** Co-corresponding Authors, Kunal Rai, Jeffrey Myers.

## Abstract

Head and neck squamous cell carcinoma (HNSCC) is a heterogeneous disease with significant morbidity and mortality and frequent recurrence. Pre-NGS efforts to transcriptionally classify HNSCC into groups of varying prognosis have identified four accepted molecular subtypes of disease: Atypical (AT), Basal (BA), Classical (CL), and Mesenchymal (MS). Here, we investigated the active enhancer landscapes of these subtypes using representative HNSCC cell lines and identified samples belonging to the AT subtype as having increased enhancer activity compared to the other 3 HNSCC subtypes. Cell lines belonging to atypical subtype were more resistant to bromodomain inhibitors (BETi). PRO-Seq experiments that both TCGA tumors and AT cell lines showed higher eRNA transcripts for enhancers controlling BETi resistance pathways, such as lipid metabolism and MAPK signaling. Additionally, HiChIP experiments suggested higher enhancer-promoter (E-P) contacts in the AT subtype, including on genes identified in the eRNA analysis. Consistently, known BETi resistance pathways were upregulated upon exposure to these inhibitors. Together, our results identify that the AT subtype of HNSCC is associated with high enhancer activity, resistance to BET inhibition, and signaling pathways that could serve as future targets for sensitizing HNSCC to BET inhibition.

## INTRODUCTION

Head and neck squamous cell carcinoma (HNSCC) is the sixth most common cancer worldwide and the predominant form of head and neck cancer [1, 2]. In the United States, there are over 60,000 new HNSCC cases and more than 13,000 HNSCC deaths per year [1]. HNSCC covers a wide variety of anatomical sites, including the oral cavity, oropharynx, hypopharynx, and larynx [3]. The prognosis for HNSCC is overall poor with a 5-year survival of approximately 50%, which has been relatively unchanged for decades [2]. This is largely attributed to factors such as late stage at initial presentation and high rates of primary tumor recurrence [4, 5]. Treatment for HNSCC involves combinations of surgery, chemotherapy, and radiotherapy, with exact treatment plans depending on tumor location and TNM stage [1, 2].

To date, studies on HNSCC have focused largely on genomic characterizations such as exome sequencing and copy number alterations. The most common alterations, such as mutations in *TP53* at 17p13 and alterations in *p16* at 9p21, have been known for decades [1, 3]. More recent comprehensive analyses of HNSCC tumors have supported these previous findings, in addition to identifying common alterations in the Notch1 pathway and cell cycle genes [4, 6, 7]. Unfortunately, very few of these studies have resulted in clinically actionable findings. There are, however, some disputed exceptions, such as the EGFR inhibitor cetuximab, which showed benefit when combined with radiotherapy [5].

One interesting result of these and other studies is the notion of molecular subtypes of disease. Inspired by similar studies in other tumors such as breast, lung, and brain, groups focused on HNSCC have used transcriptomic data from patient samples to classify head and neck tumors into 4 subtypes: Atypical (AT), Basal (BA), Classical (CL), and Mesenchymal (MS) [7–9]. These studies have largely focused on the relationship of these subtypes to genomic alterations, such as mutation patterns, copy number changes, or alterations in key transcription factor expression, and clinical features, such as progression free survival and lymph node metastasis at time of diagnosis.

While there have been several studies studying genomic associations with the four HNSCC molecular subtypes, there have been very few studies describing the epigenomic features of the subtypes. Since HNSCC subtypes are defined by their transcriptomic signatures and have distinct mutational landscapes and copy number alterations, it stands to reason they would also have unique epigenomic features, such as enhancer landscapes, that may be, in part, driving the defining transcriptomic signatures. The importance of histone modifications in HNSCC is further evidenced by the finding that global levels of certain histone tail modifications correlate with clinical measurements such as tumor stage, cancer-specific survival, and disease-free survival in oral squamous cell carcinoma [10]. Because there are currently only a sparse number of HNSCC epigenomics datasets, particularly in the realm of histone modifications and chromatin regulation, there remains an unmet need to investigate these aspects of gene regulation and leverage newly discovered biology to better define the disease and develop new therapeutic approaches [11, 12].

Through mapping of H3K27ac-marked active enhancers in 28 HNSCC cell lines, we demonstrate that AT subtype is characterized by high enhancer activity. Consistently, AT subtype was associated with resistance to enhancer-blocking bromodomain inhibitors (BETi). BETi resistance pathways specifically showed high enhancer activity as measured by nascent eRNAs and enhancer-promoter contacts providing mechanistic insights into aggressive nature of AT subtype. Overall, our data suggest high enhancer activity as an epigenetic feature of atypical HNSCCs.

## RESULTS

### HNSCC Cell Lines Assigned to Known Molecular Subtypes Have Similar Mutation Profiles and Tissue Origins

Identifying inter-tumor heterogeneity can help better understand the diversity of biological mechanisms driving the neoplastic phenotypes within pathology-based tumor types (e.g. breast cancer, colon cancer, etc.) and discover targeted therapies for specific patient populations [13–16]. We sought to define heterogeneity within HNSCC patients, specially at the epigenetic level. To this end, we first leveraged published work by Yu et al., which extensively studied CCLE cancer cell lines and their appropriateness as models of human cancer by comparing them to corresponding TCGA tumors [17]. As part of this work, the group generated “templates” of gene expression values for numerous subtypes in 9 different tumor types. Using the HNSCC templates from this study (one per molecular subtype) and RNA-seq data from the panel of cell lines available to us (**Table S1**), we utilized the nearest template prediction method, as implemented in the CMScaller R package, to assign our cell lines to their most representative molecular subtype **(Figure 1A)** [18, 19]. After selecting only samples with an assignment FDR < 0.1, twenty-eight HNSCC cell lines were successfully matched to a molecular subtype, resulting in 7 AT samples, 9 BA samples, 5 CL samples, and 7 MS samples **(Figure 1B)**. Analysis of RNA-seq data in one-versus-rest comparisons demonstrated varying levels of differential expression based on subtype, with a large number of upregulated genes (FC > 3, *n* = 756) in the AT subtype **(Figure 1C)**.

**Figure 1:**
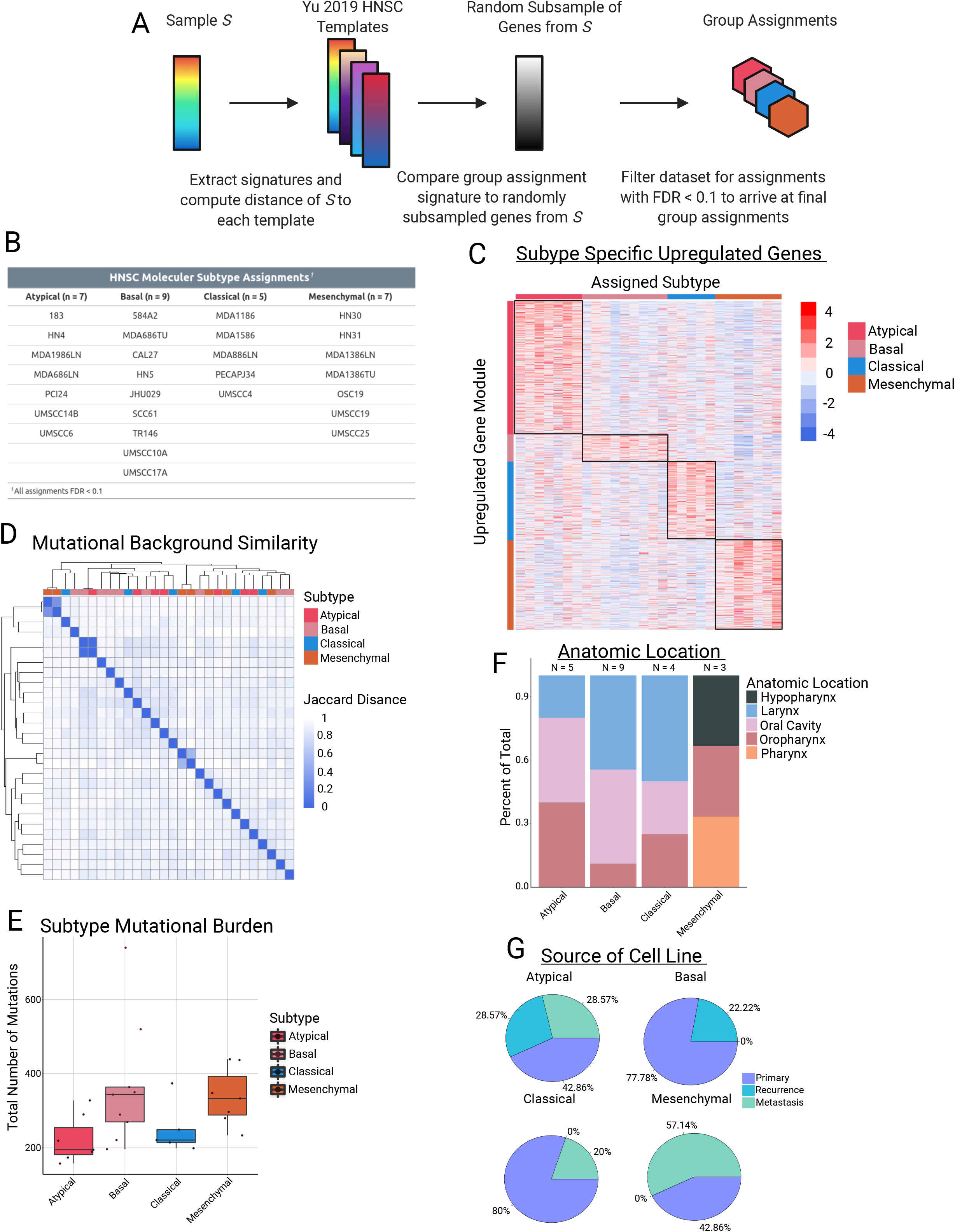
Cell line subtype assignments and characteristics. **A**, Schematic of workflow used to assign HNSC cell lines to their respective subtypes using RNA-seq data. **B**, Table of subtype assignments for each of the 28 cell lines used in this study. **C**, Heatmap of gene expression modules in each molecular subtype, defined as FC > 3 in a one-vs-rest comparison. **D**, Hierarchical clustering of the 28 cell lines based on Jaccard distance metrics obtained from binarized mutation counts from WES data. **E**, Boxplots demonstrating total number of mutations in each sample, grouped by molecular subtype (*p* = NS for each comparison). **F**, Stacked barplot showing distribution of cell line anatomic location for each molecular subtype. **G**, Pie chart showing percentage of samples in each molecular subtype that came from primary, recurrent, or metastatic lesions.

To further investigate a potential genomic basis that could be driving the transcriptomic partitioning into molecular subtypes, we investigated whole-exome sequencing (WES) data on our panel of cell lines. We first clustered our lines based on their mutational background, and, interestingly, we did not observe any clustering of samples from the same molecular subtype. In fact, the only examples of tight clusters in the data came from 3 matched pairs of cell lines in which the samples were either from a primary or metastatic lesion of the same patient **(Figure 1D)**. Similarly, we did not observe any significant differences between subtypes based on total mutational burden **(Figure 1E)**. Importantly, observed clustering was neither associated with the anatomic site of origin of the primary tumor from which each the cell lines were derived, nor with the type of tumor the sample was from (e.g. primary vs. recurrence). With the possible exception of the MS subtype, all of the HNSCC subtypes had a fairly equal distribution of samples from the oral cavity, oropharynx, and larynx **(Figure 1F)**. With respect to cell line source, the AT subtype was the only one to contain cell lines from all 3 groups of samples (primary, recurrence, and metastasis), while BA contained only primary and recurrence, and CL and MS contained only primary and metastasis **(Figure 1G)**. Taken together, these results demonstrate that HNSCC molecular subtypes can be successfully assigned to cell lines using RNA-seq data, and that, despite their transcriptomic differences, the unique HNSCC subtypes do not have significantly different mutational backgrounds, overall mutation burden, or tissues of origin from one another.

### HNSCC Molecular Subtypes are Associated with Distinct Enhancer Landscapes

Transcription of a gene is regulated by concerted action of multiple complexes on specific epigenetic elements located in *cis* or *trans* the gene promoter. Enhancers are a major component of the gene regulation circuits and known to be deregulated in cancers [20, 21]. They act as binding platforms for transcription factors that, upon various environmental cues relayed by the cell surface signaling pathways, cooperate with chromatin modifying and remodeling machinery to activate target genes [20]. We hypothesized that these transcriptional subtypes could have underlying differences in gene regulatory landscapes that explain the observed transcriptomic differences. To investigate these differences, we generated enhancer profiles for each cell line by performing ChIP-seq for the H3K27ac histone mark, which is widely used as a marker of active enhancers [22, 23]. We next generated consensus peak sets for each subtype by overlapping the enhancer regions of all cell lines within a subtype and taking the set of enhancers that occurred in 2 or more samples of the subtype. This resulted in 4 total consensus peak sets, each representing a unique subtype (i.e., one consensus peak set per subtype). We noted significantly distinct enhancer peak enrichment between the four molecular subtypes **(Figure 2A)** [24, 25].

**Figure 2:**
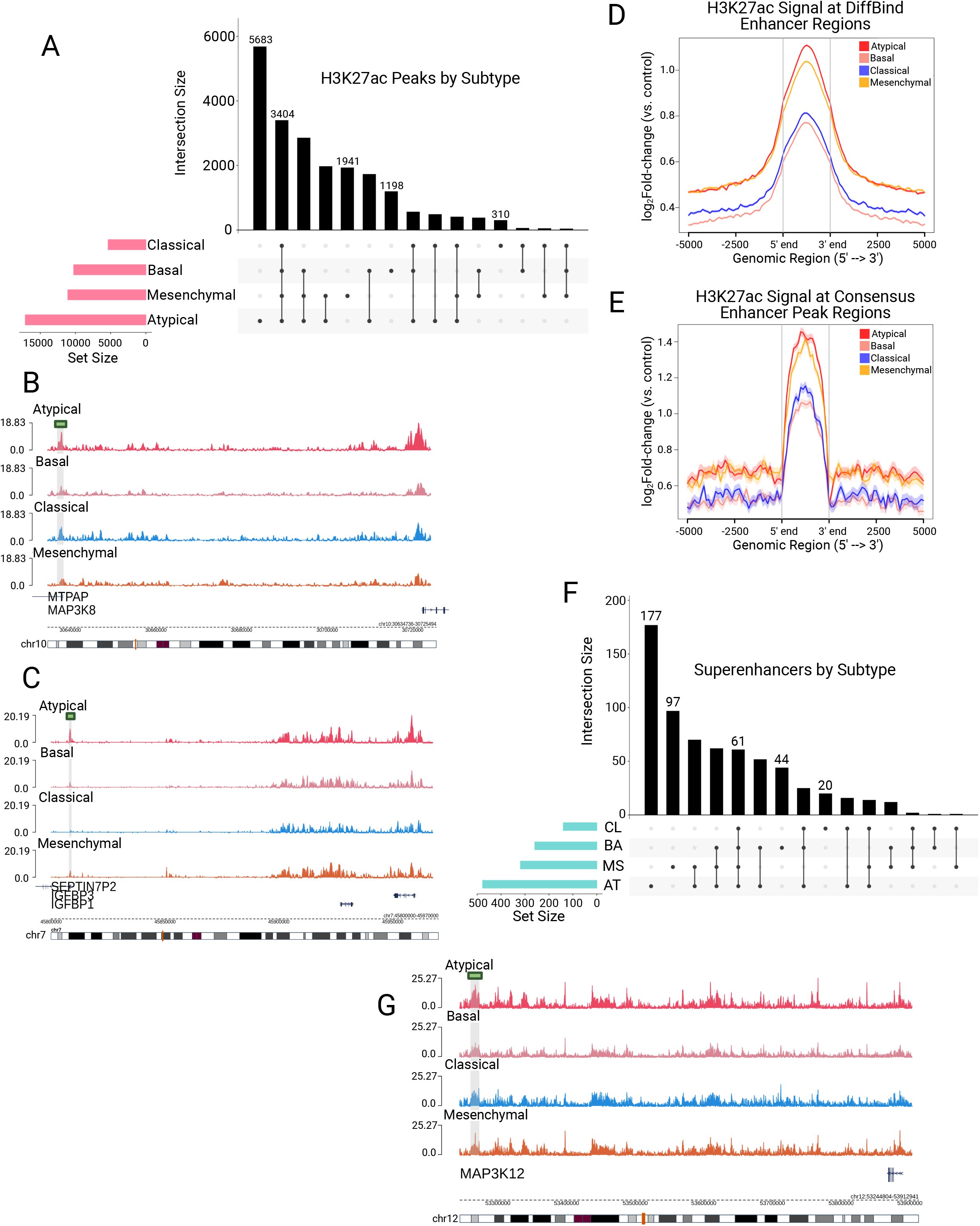
The Atypical subtype is associated with high numbers of unique enhancer peaks on genes related to lipid metabolism and MAPK signaling. **A**, UpSet plot showing the total number of H3K27ac typical enhancer peaks in each molecular subtype (pink horizontal barplot), as well as the number of peaks in each possible intersection of peaksets (black bars and dot plot). **B, C**, Visualization of mean bigWig signal for each subtype at **(b)** MAP3K8 and **(c)** IGFBP3 enhancer loci containing H3K27ac peaks unique to the AT subtype (green bar/grey shading). **D, E**, H3K27ac ChIP-seq enrichment plots of enhancer loci common to all HNSC molecular subtypes, defined as **(d)** any peak contained within 2 or more individual samples or **(e)** the 3,404 peaks shared among all consensus peaksets in **(a)**, demonstrating the strongest signal in the AT subtype. **F**, UpSet plot showing the total number of super enhancer peaks in each molecular subtype (blue horizontal barplot), as well as the number of super enhancer peaks in each possible intersection of peaksets (black bars and dot plot). **G**, Visualization of mean bigWig signal for each subtype at a MAP3K12 super enhancer containing peaks unique to the AT subtype (green bar/grey shading).

Notably, we discovered the AT subtype has a much larger set of consensus enhancers than any of the other subtypes, and the most common subset of enhancers in our analysis is the set that is unique to the AT subtype **(Figure 2A)**. To investigate the function of the 5,683 enhancers unique to the AT subtype, we utilized data from Cao et al. [26], which constructed enhancer-target networks across multiple cancer and sample types, to assign each of these enhancers to their target genes **(Figure 2B-C)**. These genes were then used for pathway enrichment analysis, which revealed enrichment for pathways involved in lipid metabolism, MYC signaling, and MAPK signaling **(Figure 2B-C, Table S2).** Notably, each of these pathways have been previously implicated as mechanisms underlying resistance to BET inhibition across many cancer types [27–31].

In addition to identifying unique enhancers, we investigated the total H3K27ac signal enrichment across enhancers shared by all 4 subtypes to determine if, in addition to the largest number of H3K27ac peaks, the AT subtype also had greater signal enrichment overall. Indeed, using two separate methods to arrive at a “shared” H3K27ac peak set, we observed that the AT subtype had more enrichment of H3K27ac signal across enhancers shared among all HNSCC subtypes **(Figure 2D-E)**. In agreement with our typical enhancer analysis, we found that the AT subtype also harbored the largest number of called superenhancers **(Figure 2F-G)**. Further, linking of these super enhancers to their gene targets not only displayed enrichment for MAPK signaling and lipid metabolism, but also identified enrichment for PI3K and WNT-β-catenin signaling **(Table S3)**. These results demonstrate that the AT subtype of HNSCC is enriched for H3K27ac-marked typical enhancers and superenhancers compared to other HNSCC subtypes which may activate important cell signaling pathways that are associated with aggressive HNSCCs.

### The Atypical Subtype Is More Resistant to BET Inhibition

Recently, the realm of “epigenetic” therapies for cancer has been of major interest for both research and in clinical applications [11, 32, 33]. Targeting epigenetic modifications and the proteins that regulate their placement and/or removal is a particularly attractive approach to cancer therapeutics since these modifications are generally considered to be reversible, particularly when compared to more “permanent” changes such as mutations and copy number alterations. One class of compounds with numerous clinical trials for a variety tumor types is BRD and extraterminal domain (BET) inhibitors, which function by inhibiting the “reader” proteins responsible for recognizing and propagating the signal of acetylated histone residues. These inhibitors have been used in the prior studies as enhancer-blocking agents. The pathways activated by AT-specific enhancers and superenhancers are also well-characterized mechanisms of BET inhibitor resistance **(Table S2-3)** [27, 34, 35]. Hence, we hypothesized that the AT subtype may be differentially responsive to BET inhibition.

We first investigated HNSCC cell line response to JQ1 using the publicly available CCLE drug response data [36–38]. We used the available CCLE RNA-seq data to assign samples to their respective HNSCC molecular subtype, then compared their response to BET inhibition in this dataset. Interestingly, we found that the AT samples have a lower JQ1 AOC (Area Over the Curve where higher values indicate resistance) than those in the non-AT group (*p* = 0.0503, Welch’s *t*-test) **(Figure 3A, S1A)**, indicating the AT subtype is more resistant to JQ1 treatment. To extend this analysis, we selected 2 compounds currently being evaluated in clinical trials, OTX015 (Birabresib) and PLX51107, and performed drug response assays in our HNSCC cell lines [32]. For each compound, we selected representative cell lines for each molecular subtype (3 AT, 3 BA, 2 CL, and 3 MS), treated with the respective compound for 72 hours, then computed the GR_AOC_ of each cell line using cell confluence as a proxy for cell number. We elected to use the GR_AOC_ metric for drug response since GR metrics have been demonstrated to be more reproducible than traditional metrics, such as Area Under the Curve (AUC) and IC_50_, when measuring drug sensitivity in cancer cell lines [39]. As we anticipated based on our previous analysis, the BET inhibitor PLX51107 demonstrated significantly lower GR_AOC_ values in the AT subtype compared to any other HNSCC subtype, indicating an increased resistance to treatment in that group **(Figure 3B)**. The inhibitor OTX015 also displayed a similar trend towards increased resistance in the AT subtype that was similar, but more pronounced than, to the JQ1 response data in the CCLE database **(Figure S1A-B)**.

**Figure 3:**
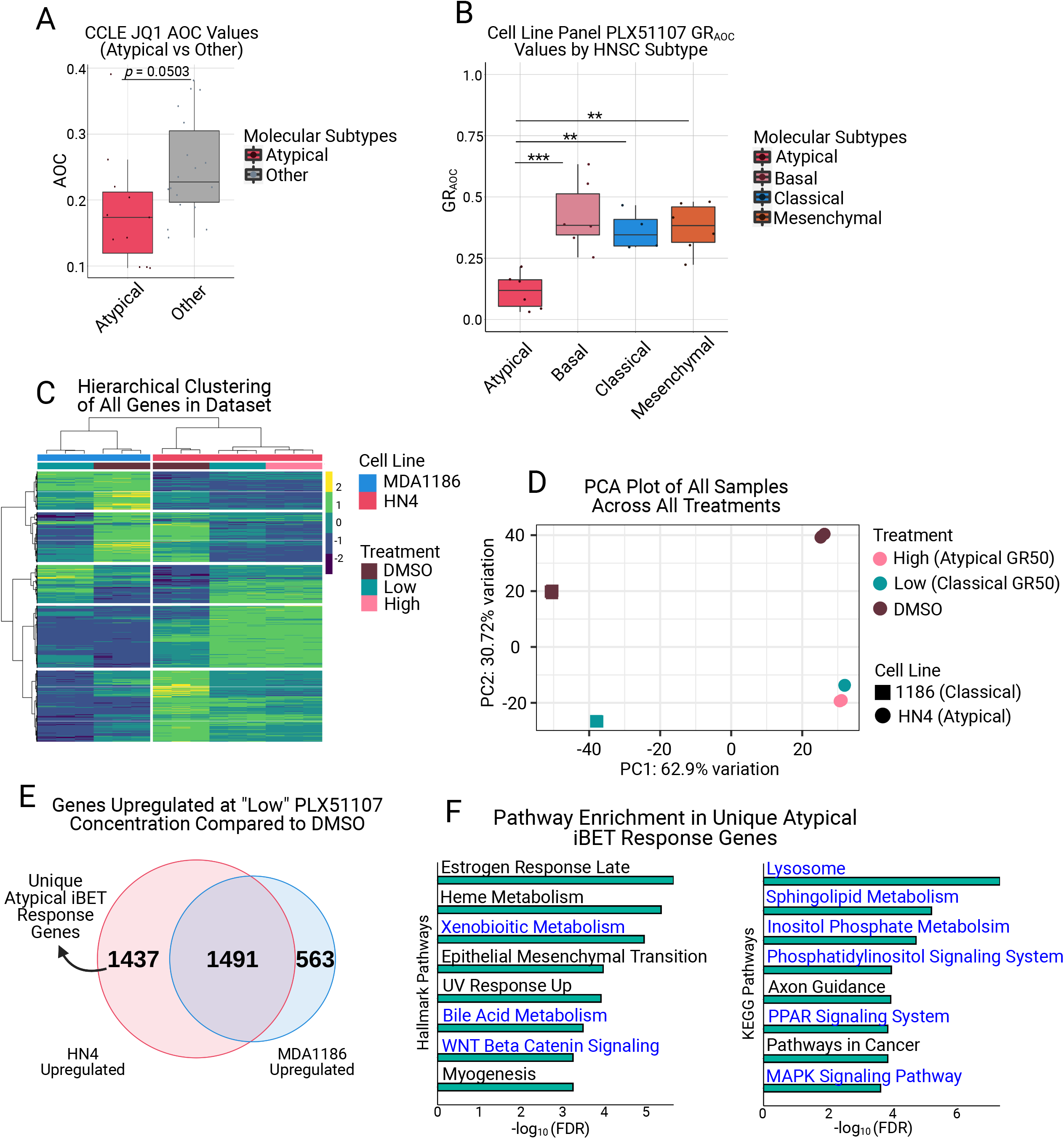
Atypical HNSC shows increased resistance to BET inhibition and uniquely upregulates genes associated with resistance pathways upon treatment. **A**, Atypical samples in the HNSC CCLE dataset demonstrate lower JQ1 AOC values than non-Atypical samples (*p* = 0.0503, Welch’s t-test). **B**, Drug response assays with the BET inhibitor PLX51107 demonstrate the Atypical subtype is significantly more resistant to BET inhibition than other molecular subtypes (*** *adj.p* < 0.001, ** *adj.p* < 0.01). **C**, Hierarchical clustering of all genes from HN4 and MDA1186 samples treated with DMSO, PLX51107 at GR_50_ MDA1186 (low), or GR_50_ HN4 (high). **D**, PCA plot of samples as described in **(c)**, displaying separation on the basis of cell line (PC1) and treatment status (PC2). **E**, Overlap of genes upregulated (|log_2_fold-change| > 1.5 & FDR < 0.05) in HN4 and MDA1186 at PLX51107 low concentration; numbers in venn diagram represent size of set. **F**, Horizontal barplots of Hallmark (left) and KEGG (right) pathway enrichment results from the 1,437 genes uniquely upregulated by HN4 in **(e)**; pathways highlighted in blue are associated with known mechanisms of BET inhibitor resistance.

To better understand the mechanisms behind this observed resistance to treatment, we selected one representative cell line from the AT subtype (HN4) and one cell line from the non-AT subtypes (MDA1186, Classical subtype), treated with PLX51107 or DMSO, and performed gene expression analysis using mRNA-seq profiling in each condition. MDA1186 was treated with PLX51107 at its own GR_50_ value (hereafter referred to as “low” concentration), and HN4 was treated at its own GR_50_ value (hereafter referred to as “high”), as well as the low concentration. To ensure PLX51107 behaved similarly to other published BET inhibitors, we created BET inhibitor response signatures using publicly available data and, using GSEA, confirmed that the response of the AT and non-AT cell lines to PLX51107 was consistent with previously documented BET inhibitor response signatures **(Figure S1C-D)** [40, 41]. Hierarchical clustering of all genes in the RNA-seq dataset demonstrated 2 major clusters, one per cell line, as well as 2 sub-clusters, either DMSO or PLX51107-treated, per major cluster **(Figure 3C)**. To further examine the differences in response to drug treatment, we performed a principal component analysis (PCA), which revealed a first component driven by the cell line identity, and a second component driven by treatment status **(Figure 3D)**. Examining the results from the hierarchical clustering and PCA analysis together, we noted that the majority of transcriptional response to BET inhibition in the AT cell line occurs at the lower drug concentration, and only a minority of gene expression changes occur between the low and high concentrations **(Figure 3C-D)**. Closer inspection of the gene expression heatmap indicates that the genes that are specifically upregulated in the AT subtype after PLX51107 treatment, but not in the non-AT subtypes, may be responsible for mediating resistance to BET inhibition **(Figure 3C)**.

To investigate these uniquely upregulated genes, we overlapped the set of genes upregulated by the AT subtype and by the non-AT subtype at the low PLX51107 concentration. As we expected, we discovered a set of 1437 genes uniquely upregulated in the AT subtype after BET inhibition **(Figure 3C, E)**. Pathway analysis of these 1437 genes using the Hallmarks and KEGG gene sets reveals enrichment for multiple pathways previously demonstrated to convey resistance to BET inhibition and identified by our previous enhancer-based analysis, including MAPK signaling, WNT-β-catenin signaling, phosphatidylinositol signaling, and lipid metabolism pathways **(Figure 3F)** [27–31, 34, 35].

These results support our previous hypothesis that the AT subtype is more resistant to BET inhibition than other HNSCC subtypes and suggest the enrichment of H3K27ac-marked enhancers involved in these pathways is a contributing factor to the observed resistance.

### BET Inhibitor Resistance Is Mediated by Baseline Enhancer Activity and Chromatin Structure

Our previous analyses have indicated that the AT subtype of HNSCC has two intriguing properties with respect to BET inhibition: first, H3K27ac-marked enhancers unique to the AT subtype regulate genes enriched for known BET inhibitor resistance pathways, and, second, the AT subtype is able to uniquely upregulate genes enriched for BET inhibitor resistance pathways after BET inhibitor treatment **(Figure 2B-C, G; Table S2-3; Figure 3E-F)**. Because of these observations, we suspected the AT subtype may have a stronger baseline enhancer activity at genes involved in resistance pathways and that these genes may have higher numbers of enhancers-promoter contacts, enabling a more robust response to BET inhibitor treatment.

To investigate the activity of enhancers involved in regulating baseline resistance gene expression, we performed precision nuclear run-on sequencing (PRO-seq) to investigate the enhancer RNA (eRNA) landscape of the genes [42]. Enhancer RNAs are a recently discovered class of non-coding transcripts found at active enhancers that arise from the transcription of the enhancer elements themselves and are involved in functions such as regulating gene transcription and controlling enhancer-promoter looping [43–45]. We used PRO-seq, with a particular focus on eRNAs, to investigate differential enhancer activity in our AT subtype. For this experiment, we expanded our AT group to include the 3 cell lines from our drug assay, and we expanded the non-AT group to include the 2 cell lines from the classical subtype used in our drug assay. Differential expression analysis of PRO-seq-defined eRNAs revealed 321 differentially expressed eRNAs, with 207 being upregulated and 114 being downregulated **(Figure 4A)**.

**Figure 4:**
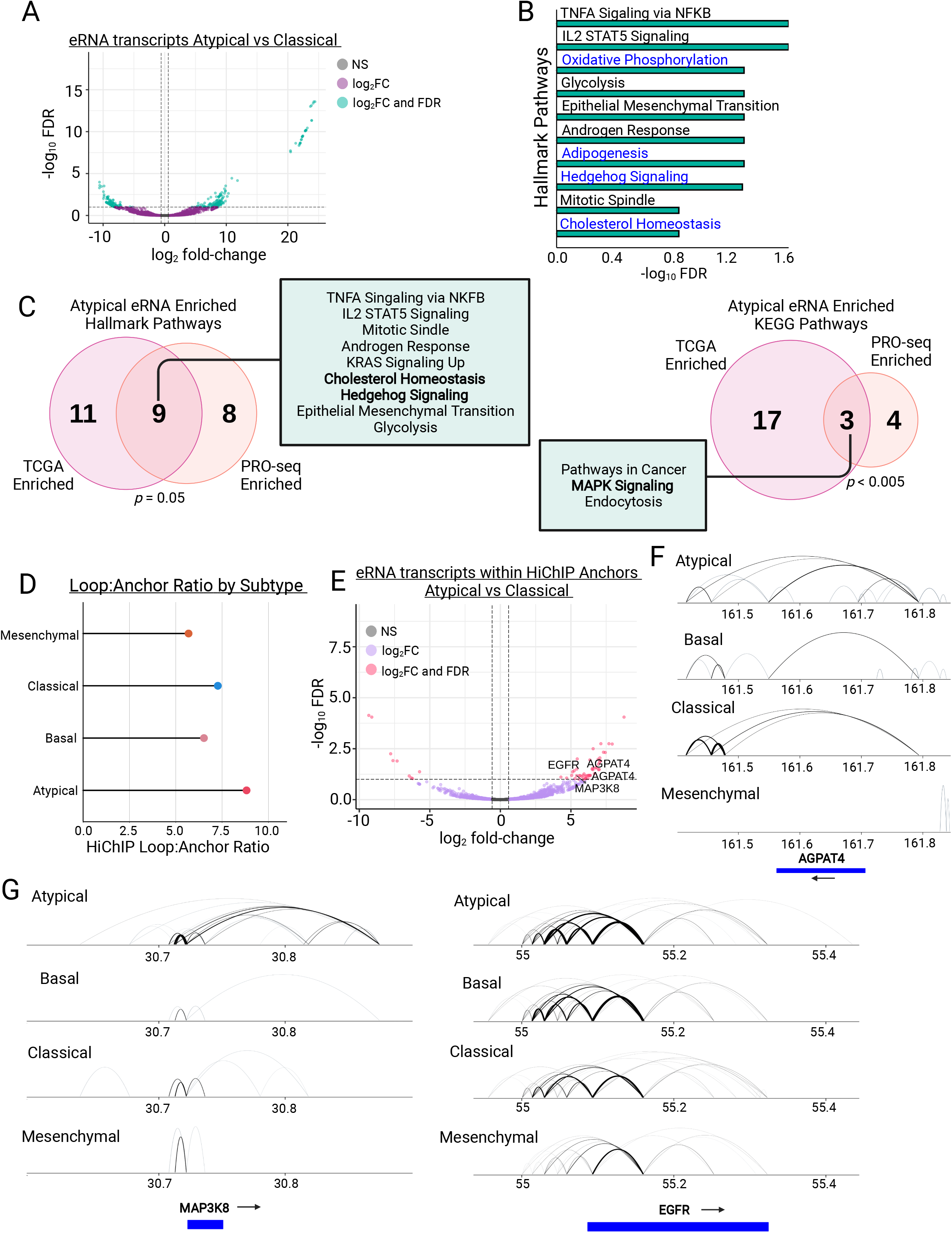
Enhancers of MAPK signaling, WNT signaling, and Cholesterol Homeostasis genes display increased eRNA transcription and enhancer-promoter looping in Atypical HNSC. **A**, Differential transcription (|log_2_fold-change| > 1.5 & FDR < 0.1) of eRNAs between the Atypical and Classical subtypes as measured by PRO-seq (green dots meet fold-change and FDR thresholds, purple dots meet fold-change threshold only). **B**, Hallmark pathway enrichment analysis of genes linked to eRNAs with significantly increased transcription from **(a)**; pathways in blue have been previously associated with BET inhibitor resistance. **C**, Overlap of hallmark (left) and KEGG (right) pathway enrichment results between PRO-seq-determined significantly enriched eRNAs from **(a)** and **(b)** and TCGA-measured differentially expressed eRNAs between Atypical and non-Atypical samples; *p* values represent hypergeometric tests of gene set enrichment result overlaps; bolded terms represent shared pathways associated with BET inhibitor resistance. **D**, Lollipop plot demonstrating the loop count:anchor count ratio of H3K27ac HiChIP data for each molecular subtype. **E**, Volcano plot of differentially transcribed (|log_2_fold-change| > 1.5 & FDR < 0.1) eRNAs between the Atypical and Classical subtypes after filtering transcripts for only those contained within H3K27ac HiChIP anchors (pink dots meet foldchange and FDR thresholds, purple dots meet fold-change threshold only). **F**, Joined Hallmark and KEGG pathway enrichment analysis of genes linked to differentially transcribed eRNAs in **(e)**; pathways in blue have been previously associated with BET inhibitor resistance. **G**, Visualization of H3K27ac HiChIP loops at the MAP3K8 locus (left) and EGFR locus (right) in all 4 HNSC molecular subtypes.

To assess the likely functional output of these eRNAs, we assigned each eRNA to its nearest gene and performed pathway enrichment analysis, which demonstrated an enrichment in multiple metabolic pathways, including lipid metabolism and cholesterol homeostasis, and hedgehog signaling **(Figure 4B)**. These findings are largely in agreement with our previous enhancer-based analysis of H3K27ac-linked genes, which displayed enrichment for similar BET inhibitor resistance associated pathways **(Table S2)**. To extend this finding to human tumors, we leveraged data from recent publications investigating eRNA expression in TCGA tumors [33, 46]. After assigning all the HNSCC TCGA tumor samples to their molecular subtype, we examined eRNA expression in the AT subtype compared to non-AT samples and found the AT subtype upregulated 1,867 eRNAs. After linking these eRNAs to their nearest gene, we found that, in agreement with our PRO-seq data from HNSCC cell lines, these genes were enriched for cholesterol homeostasis, hedgehog signaling, and MAPK signaling function **(Figure 4C)**. These results support the hypothesis that the AT subtype has active enhancers, as measured by eRNA expression, enriched for signaling pathways that have been demonstrated to confer resistance to BET inhibition across cancer types.

To assess the enhancer-promoter contacts in our HNSCC cell lines, we performed HiChIP for H3K27ac-marked histones to capture the enhancer-promoter looping involving this enhancer mark [47]. Consistent with enhancer analyses, we discovered that the AT subtype has the highest ratio of H3K27ac-mediated loops to H3K27ac anchors across all 4 HNSCC subtypes, indicating the AT subtype may have more redundancy in its enhancer architecture than the other subtypes **(Figure 4D)**. To assess if these loops are related to enhancer function, we overlapped our PRO-seq called eRNA enhancer regions with the H3K27ac HiChIP anchor data and performed differential expression analysis of this subset of eRNAs. Pathway analysis for the upregulated eRNAs contained within H3K27ac HiChIP anchors (64 upregulated eRNAs) demonstrated results consistent with our previous analyses with enrichment for MAPK signaling, WNT-β-catenin signaling, and cholesterol homeostasis **(Figure 4E-F)**. Further, inspection of genes identified by this integrative analysis revealed increased number of contacts between respective gene promoters and H3K27ac-marked enhancers, supporting the association of eRNA expression with active enhancers and enhancer-promoter loop formation **(Figure 4G-H)**.

Overall, insights from the eRNA expression and HiChIP data support a model in which the AT subtype has more active enhancers regulating genes associated with lipid metabolism and MAPK signaling, and AT enhancers have, on average, a higher level of redundancy in their control of gene expression than non-AT enhancers by forming larger number of enhancer-promoter contacts.

## DISCUSSION

Here, we have demonstrated that HNSCC molecular subtypes have largely similar mutational backgrounds, mutational burden, and tissues of origin. In contrast, the enhancer landscapes, marked by histone H3K27 acetylation, are distinct among subtypes. In particular, we discovered the Atypical subtype has the highest number of enhancer and super enhancers, as well as the most enhancer signal at common enhancer regions, and a global increase in enhancer-promoter loop formation. We also demonstrate that the Atypical subtype is more resistant to BET inhibition and that, upon treatment with BET inhibitors, the Atypical subtype is able to uniquely upregulate genes associated with cell growth and BET inhibitor resistance pathways (MAPK signaling, WNT signaling, and lipid metabolism) [27–31, 34, 35]. Further, we demonstrate a significant baseline upregulation of eRNA transcription from the enhancers of genes involved in BET inhibitor resistance pathways such as lipid metabolism and hedgehog signaling [27] in the Atypical subtype. Interestingly, many of these genes with increased eRNA expression in their enhancers were also found to have baseline increased enhancer-promoter looping. Together, our findings suggest that the Atypical subtype of HNSCC is characterized by high enhancer activity, which likely drives the expression of pathways known to confer resistance to BET inhibition.

Delineation of HNSCC in 4 subtypes was originally proposed by Walter et al. and the TCGA HNSCCC study [9, 48]. These two manuscripts largely focus on genomic alterations, such as copy number alterations and somatic mutations, and only one epigenetic element, in the form of DNA methylation, was assessed in the TCGA paper. As such, limited epigenomic data for HNSCC is available [12]. Despite these limitations, interest in therapies that target the epigenome continues to grow, indicating a need for more studies that focus on the epigenome of HNSCC [32, 49]. The work presented here is, to our knowledge, the first to characterize the enhancer landscape of HNSCC based on the Walter/TCGA molecular subtypes. Interestingly, we identify HPV-negative samples belonging to Atypical subtype, which has traditionally been associated with HPV-positive or “HPV-like” samples, have increased enhancer activity compared to the non-AT subtypes. This activity is measured by increased H3K27ac peak counts, increased H3K27ac signal at common enhancer peaks, global increases in enhancer-promoter looping, and significant upregulation of eRNA expression compared to non-AT samples. This finding suggests that defining features of the Atypical subtype are enhancer architecture and activity - two key epigenomic aspects of HNSCC subtypes that were not previously explored. Clinical and translational significance of enhancer-based classification was shown by our recent studies in other tumor types like colorectal cancer [50] and MPNST [51].

Unfortunately, BET inhibitors have shown limited promise in clinical studies in solid tumors [52]. However, in specific solid tumor contexts, such as BRD4-NUT midline carcinoma, BET inhibitors have had very encouraging results in clinical trials [52–54]. Considering these findings, identifying subsets of patients with tumor biology favorable or unfavorable to BET inhibitor response could improve the clinical utility of these compounds. Since BET inhibitors inherently rely on modulating the reader protein of H3K27ac-marked enhancers in target cells, understanding enhancer landscapes and their role in BET inhibitor response becomes an important first step in sorting patients into “favorable” or “unfavorable” groups [33, 55]. In our work, we discovered that the Atypical subtype is significantly more resistant to BET inhibition than other HNSCC subtypes, and that this resistance seems, at least in part, mediated by increased enhancer activity on pathways associated with lipid and cholesterol metabolism, MAPK signaling, and WNT-β-catenin signaling. Accordingly, we expect that including compounds that target these pathways in combination with BET inhibitors may sensitize otherwise resistant tumors to BET inhibition and expand the current chemotherapeutic repertoire for HNSCC treatment. As such, other enhancer/transcription blocking inhibitors, such as those against CDK9 [56], could be tested in such enhancer-based subtypes.

While our work focused on HPV-negative HNSCC, our findings suggest increased enhancer activity on genes involved in lipid and cholesterol metabolism, MAPK signaling, and WNT-β-catenin signaling may serve as a general mechanism of baseline resistance to BET inhibition. Since enhancer architecture is a critical component of cell identity, it is possible that, moving forward, an assessment of a tumor’s baseline enhancer activity could serve as a potential epigenomic biomarker of response to BET inhibition and aid in tailoring treatment in a patientspecific manner [57, 58]. This could be especially useful in the case of HNSCC, where subtype-specific and tumor-specific treatments are generally lacking [49].

## METHODS

### Cell culture

Human HNSCC cell lines were acquired and characterized as previously described [59]. Briefly, cell lines were cultured in DMEM supplemented with 10% FBS, L-glutamine, sodium pyruvate, nonessential amino acids, vitamins, and 1% penicillin-streptomycin. All cell lines were cultured at 37 °C in an atmosphere of 5% CO_2_.

### RNA-sequencing processing and analysis

RNA-seq data for cell line subtype assignments were obtained as raw counts, processed as previously described [60, 61]. To assign HNSCC cell lines to their representative subtypes, we used the HNSCC gene list templates generated by Yu et al. [17] and utilized the CMScaller workflow and implementation of the NTP algorithm to find the closest matching subtype for each cell line (FDR < 0.1) based on their transcriptomic profiles [18, 19]. Upregulated genes for each subtype were computed using CMScaller in a one-vs-rest fashion.

For the PLX51107 treatment RNA-seq experiments, representative cell lines were selected for the Atypical subtype (HN4) and a non-Atypical subtype (MDA1186, Classical subtype) and were treated with DMSO, GR_50_ of the MDA1186, or GR_50_ of HN4 for 24 hours prior to RNA isolation. RNA extraction was performed using an RNeasy Mini Kit per manufacturer’s instructions (Qiagen). Isolation of mRNA was performed using NEBNext Poly(A) mRNA Magnetic Isolation Module and libraries were prepared using the NEBNext Ultra II Directional RNA Library Prep kit (New England BioLabs). Library quality was checked on an Agilent TapeStation 4150 and quantified by Qubit 2000 fluorometer (Invitrogen). Libraries were pooled in equimolar ratios and sequenced on Illumina NovaSeq6000 SP runs with paired-end 100-bp reads at the The Advanced Technology Genomics Core (ATGC) at MD Anderson Cancer Center.

PLX51107 treatment RNA-seq raw reads were processed using the provided pipeline: https://github.com/sccallahan/QUACKERS_RNAseq-pipeline. In brief, raw reads were aligned to the hg19 genome using STAR v2.7.2b [62] and quality checked using FastQC v0.11.8 [63] (https://www.bioinformatics.babraham.ac.uk/projects/fastqc/). Counts were generated using featureCounts from subread v1.6.3 80 [64]. Downstream normalization and differential expression analysis were performed using DESeq2, with size factors being calculated using data-driven housekeeping gene method as implemented in the CustomSelection R package [65, 66]. Pathway enrichment analyses were performed using GSEA’s pre-ranked list option [67]. Overlaps of HN4 and MDA1186 low dose PLX51107 differentially expressed genes were performed using the VennDiagram package in R, and the HN4 uniquely upregulated gene list was subjected to pathway enrichment analysis using the gsea-msigdb online tool (http://www.gsea-msigdb.org/gsea/msigdb/annotate.jsp).

CCLE RNA-seq data were downloaded as raw counts from the depmap download portal (https://depmap.org/portal/download/). Subtype assignment and downstream analysis were carried out as above.

### Whole exome sequencing processing and analysis

WES data was processed as previously described and obtained as a MAF file from the authors [60, 61]. To cluster the cell lines based on mutation background, all mutation calls were binzarized to 1 or 0 to represent “mutated” or “not mutated,” respectively. The Jaccard distance matrix was then computed, and the resulting matrix was clustered using Ward’s minimum variance method. Total mutational burden was calculated by summing the number of mutations per sample, then grouping the samples based on their assigned molecular subtype. Data for cell line tissue of origin and “source” were obtained as previously described [59].

### ChIP-sequencing processing and analysis

ChIP assays were performed as described previously [68]. Briefly, approximately 2 × 10^7^ cells were harvested by scraping. Samples were cross-linked with 1% (wt/ vol) formaldehyde for 10 min at 37 °C with shaking. After quenching with 150 mM glycine for 5 min at 37 °C with shaking, cells were washed twice with ice-cold PBS and frozen at −80 °C for further processing. Crosslinked pellets were thawed and lysed on ice for 30 min in ChIP harvest buffer (12 mM Tris-Cl, 1 × PBS, 6 mM EDTA, 0.5% SDS) with protease inhibitors (Sigma). Lysed cells were sonicated with a Bioruptor (Diagenode) to obtain chromatin fragments (~200–500 bp) and centrifuged at 15,000 × g for 15 min to obtain a soluble chromatin fraction. In parallel with cellular lysis and sonication, antibodies (5 μg/3 × 10^6^ cells) were coupled with 30 μl of magnetic protein G beads in binding/blocking buffer (PBS + 0.1% Tween + 0.2% BSA) for 2h at 4 °C with rotation. The antibody used for ChIP was anti-H3K27ac (Abcam; ab4729). Soluble chromatin was diluted five times using ChIP dilution buffer (10 mM Tris-Cl, 140 mM NaCl, 0.1% DOC, 1% Triton X, 1 mM EDTA) with protease inhibitors and added to the antibody-coupled beads with rotation at 4 °C overnight. After washing, samples were treated with elution buffer (10 mM T ris-Cl, pH 8.0, 5 mM EDTA, 300 mM NaCl, 0.5% SDS), RNase A, and Proteinase K, and cross-links were reversed overnight at 37. Immune complexes were then washed five times with cold RIPA buffer (10mM Tris–HCl, pH 8.0, 1mM EDTA, pH 8.0, 140mM NaCl, 1% Triton X-100, 0.1% SDS, 0.1% DOC), twice with cold high-salt RIPA buffer (10mM Tris–HCl, pH 8.0, 1mM EDTA, pH 8.0, 500mM NaCl, 1% Triton X-100, 0.1% SDS, 0.1% DOC), and twice with cold LiCl buffer (10mM Tris–HCl, pH 8.0, 1mM EDTA, pH 8.0, 250mM LiCl, 0.5% NP-40, 0.5% DOC). ChIP DNA was purified using SPRI beads (Beckman Coulter) and quantified using the Qubit 2000 (Invitrogen) and TapeStation 4150 (Agilent). Libraries for Illumina sequencing were generated following the New England BioLabs (NEB) Next Ultra DNA Library Prep Kit protocol. Amplified ChIP DNA was purified using double-sided SPRI to retain fragments (~200–500 bp) and quantified using the Qubit 2000 and TapeStation 4150 before multiplexing.

Raw fastq reads for all ChIP-seq experiments were processed using a snakemake based pipeline https://github.com/crazyhottommy/pyflow-ChIPseq. Briefly, raw reads were first processed using FastQC [63] (https://www.bioinformatics.babraham.ac.uk/projects/fastqc/) and uniquely mapped reads were aligned to the hg19 reference genome using Bowtie version 1.1.2 [69]. Duplicate reads were removed using SAMBLASTER [70] before compression to bam files. To directly compare ChIP-seq samples, uniquely mapped reads for each mark were downsampled per condition to 15 million, sorted, and indexed using samtools version 1.5 [71]. To visualize ChIP-seq libraries on the IGV genome browser, we used deepTools version 2.4.0 [72] to generate bigWig files by scaling the bam files to reads per kilobase per million (RPKM) and WiggleTools [73] to create average profile plots for each molecular subtype.

Peak overlaps were performed by first generating consensus peak files for each subtype, defined as any peak found in at least 2 samples from the subtype. The resulting 4 peaksets (one per subtype) were then used as input for intervene’s [25] upset module to generate upset plots and common/unique peaksets for further analysis. Gene linkage was performed using previously published enhancer-promoter linkage data from Cao et al. [26], and the resulting gene list was used as input for pathway enrichment analysis using the gsea-msigdb online tool (http://www.gsea-msigdb.org/gsea/msigdb/annotate.jsp). Enrichment plots were generated using two definitions of common peaks. The first method uses DiffBind [74] (https://bioconductor.org/packages/release/bioc/html/DiffBind.html) to define a peakset using any peak found in at least 2 samples, irrespective of subtype. The second uses the peaks found in all subtypes when overlapped using the intervene package mentioned previously. Both resulting peaksets were used as input for ngs.plot [75] to generate figures.

### Drug response assays

Cell confluence and proliferation were measured using the IncuCyte ZOOM system (Essen Biosciences). For each cell line, seeding density was optimized such that the cells would be in their exponential growth phase for the duration of drug treatment. On day 0, cells were seeded into 96 well plates and left in the incubator overnight. On day 1, media containing either drug (PLX51107 or OTX015) or DMSO was added to the wells. Plates were then left in the IncuCyte ZOOM with treated media for 72 hours, at which point cell confluence was measured. Each assay was performed in biological duplicate with technical triplicate wells. Drug response metrics were calculated using GR metrics [39] using cell confluence as a proxy for growth rate, and GR_AOC_ values from each cell line were combined based on their molecular subtype for statistical analysis.

CCLE drug response data were downloaded and processed using the PharmacoGx “auc_recomputed” dataset [76]. Compounds which were missing data for more than 25% of samples were excluded. For the JQ1 analysis, 1 sample was missing data and was imputed using predictive mean matching (as implemented in the mice package [77] (https://www.jstatsoft.org/article/view/v045i03)) on the complete, filtered drug response matrix.

### PRO-seq processing and eRNA analysis

Extraction of nuclei and precision run-on reaction was carried out as described previously [42]. Nuclei were isolated from approximately 10 million cells after treating with 12ml of ice cold swelling buffer (10mM Tris-HCl pH 7.5, 2mM MgCl2, 3mM CaCl2) for 10 minutes and scraping out the cells. After spinning at 600g for 10 minutes at 4C, supernatant was removed and cells were lysed in 10 ml of lysis buffer (10mM Tris-HCl pH 7.5, 2mM MgCl2, 3mM CaCl2, 10% glycerol, 0.5% NP-40, 4U/ml SUPERase inhibitor) on ice for 5 minutes. The lysate was spun at 600g for 8 minutes and nuclei were collected. The nuclei were then resuspended in 1ml of freezing buffer (50mM Tris-HCl pH 8.0, 5mM MgCl2, 0.1mM EDTA, 40% glycerol) and spun at 900g for 10 minutes. For performing precision nuclear run-on reaction, nuclei were resuspended in 100l of freezing buffer and added to 100l of NRO-reaction mix - NRO-reaction buffer (10mM Tris-HCl pH 8.0, 5mM MgCl2, 300mM KCl), 1mM DTT, 100U/ml SUPERase-In, 1% Sarkosyl, 250M ATP, 250M GTP, 50M biotin-11-UTP, 50M biotin-11-CTP. Reaction was carried out at 29C for 4 minutes. RNA was extracted using TRIzol. Base hydrolysis was carried out by heat denaturing briefly at 65C for 40 seconds following by cooling on ice and treatment with 1N NaOH for 6 minutes on ice. The sample with fragmented RNA was neutralized with 1M Tris-HCl pH 6.8 and isolated by passing through P-30 column (Biorad, #732-6250). The NRO-reaction products containing biotinylated RNA was purified using Streptavidin C1 beads which were washed thrice with wash buffer (0.1N NaOH, 50mM NaCl) and twice with 100 mM NaCl. The washed beads were resuspended in binding buffer (10mM Tris-HCl pH 7.4, 150 mM NaCl, 0.1% Triton X-100) and added to the sample and incubated at room temperature for 30 minutes on a rotator. After removing the supernatant using a magnetic stand, beads were washed twice with high salt wash buffer (50mM Tris-HCl pH 7.4, 2M NaCl, 0.5% Triton X-100), once with low salt wash buffer (10mM Tris-HCl pH 7.4, 300 mM NaCl, 0.1% Triton X-100) and twice with no salt wash buffer (5mM Tris-HCl pH 7.4, 0.1% Triton X-100). Beads were then resuspended in TRIzol and RNA was extracted. The bead purification of biotinylated RNA was performed once more and RNA was extracted using TRIzol to improve the purity of sample.

Libraries were generated based on previously described protocol [78]. isolated RNA samples were dephosphorylated by FastAP (ThermoFisher) and T4 Polynucleotide kinase (NEB). Samples were cleaned up using MyONE Silane beads and RNA was isolated with RLT buffer (Qiagen). To the eluted RNA, a barcoded RNA adapter (RiL 19) was ligated to the 3’ end using T4 RNA ligase (NEB). The 3’ adaptor ligated RNA was again cleaned up as mentioned above. RNA was then reverse transcribed with AR17 primer and AffinityScript reverse transcriptase (Agilent). cDNA was then cleaned up by treating with ExoSAP-IT (Affymetrix) to remove excess oligonucleotides. Excess RNA was removed from cDNA by treating with 1M NaOH at 70C for 12 minutes and neutralizing with 1M HCl. cDNA was then cleaned up with MyONE Silane beads and RLT buffer and eluted in 5mM Tris-Cl, pH 7.5. A second 5’adaptor (rand3Tr3) was ligated to cDNA with T4 RNA ligase in an overnight reaction at room temperature. The adaptor ligated cDNA was then cleaned up with MyONE Silane beads and RLT buffer and eluted in 10mM Tris-Cl, pH 7.5. cDNA samples were then PCR amplified using NEBNext^®^ Ultra^™^ II Q5^®^ Master Mix multiplexing was done with D50X and D70X primers. Libraries were size selected and purified using SPRI beads. Final libraries were quantified using D1000 tapestation and Qubit^™^ dsDNA HS Assay Kit (Thermo Fisher Scientific) and sequenced using NovaSeq6000 with 100nt paired-end format.

Fastq files from PRO-seq experiments were processed using the previously described PEPPRO pipeline [79]. Briefly, fastq files first undergo pre-processing steps of adapter removal, read deduplication, read trimming, and reverse complementation. The resulting files are then “prealigned” to the human rDNA genome to siphon off these unwanted reads. The rDNA-removed files are then aligned to the human hg19 genome using bowtie2 [80]. After quality control assessment, 5 samples from the Atypical subtype (representing the 3 unique cell lines used in the drug response assays) and 3 samples from the Classical subtype (representing the 2 unique cell lines used in the drug response assays) were carried forward for further analysis. The aligned, sorted bam files for these samples were used as input for downstream analysis using the previously described NRSA downstream analysis pipeline [81]. In brief, NRSA uses bidirectional transcription in intergenic regions to identify and call enhancers/eRNAs. A raw counts table for these eRNAs is then generated and fed into the DESeq2 tool for differential expression analysis. Identified enhancers are assigned to their nearest genes to generate a list of genes with upregulated eRNA expression in the Atypical subtype, which was then used as input for pathway enrichment analysis.

TCGA RPKM expression levels of numerous eRNAs from typical enhancers were downloaded from publicly available datasets (https://bioinformatics.mdanderson.org/Supplements/Super_Enhancer/TCEA_website/parts/3_eRNA_quantification.html) based on previously published work [46, 82]. TCGA mRNA-seq for subtype assignment was downloaded from FireBrowse (http://firebrowse.org/). TCGA samples were assigned to molecular subtypes by first generating templates based on previously assigned molecular subtypes from the initial HNSCC TCGA cohort [7]. These assigned subtypes were then expanded to the current cohort of samples by using the CMSCaller functionality described above. As before, samples were only retained for further analysis if they possessed an assignment FDR < 0.1. Samples were then grouped into “Atypical” or “Other” based on their molecular subtype, and significant differential expression of eRNAs was determined by > 1.5 fold-change in expression and FDR < 0.05. As with the PRO-seq data, these eRNAs were linked to their nearest gene using bedtools via the bedr R package [83], and the resulting gene list was used as input for pathway enrichment analysis. Intersections of the TCGA eRNA enriched pathways and PRO-seq eRNA enriched pathways were performed using the Venn Diagram package in R and significance was calculated using the hypergeometric overlap method, with a universe size set to the number of unique pathways in a particular gene set. Only pathways with an p.adjust < 0.25 with a maximum of 20 enriched pathways per gene set were included in the analysis.

### HiChIP protocol, processing, and analysis

HiChIP was performed as described [47]. Briefly, 1 × 10^7^ cells for each HNSCC cell line (1 unique cell line per HNSCC subtype) were crosslinked. In situ contacts were generated in isolated and pelleted nuclei by DNA digestion with MboI restriction enzyme, followed by biotinylation of digested DNA fragments with biotin–dATP, dCTP, dGTP, and dTTP. DNA was then sheared with Bioruptor (Diagenode); chromatin immunoprecipitation was done for H3K27Ac with use of anti-H3K27ac antibody. After reverse-crosslinking, 150 ng of eluted DNA was taken for biotin capture with Streptavidin C1 beads followed by transposition with Tn5. In addition, transposed DNA was used for library preparation with Nextera Ad1_noMX, Nextera Ad2.X primers, and Phusion HF 2XPCR master mix. The following PCR program was performed: 72°C for 5 min, 98°C for 1 min, then 11 cycles at 98°C for 15s, 63°C for 30s, and 72°C for 1 min. Afterward, libraries were twosided size selected with AMPure XP beads. Libraries were paired-end sequenced with reading lengths of 76 nucleotides.

Using HiC-Pro [84], HiChIP paired-end reads were aligned to the hg19 genome with duplicate reads removed, assigned to MboI restriction fragments, and filtered for valid interactions. Interaction matrices were then generated with the same software. To generate anchor points for downstream looping analysis, outputs from HiC-Pro were used as inputs for peak calling in HiChIP-Peaks [85]. To ensure loops were called from similar background enhancers, peaks from HiChIP-Peaks were concatenated into a single file and used as anchor point inputs for loop calling via hichipper [86]. HiChIP loop visualization was performed using DNAlandscapeR (https://molpath.shinyapps.io/dnalandscaper/).

### Statistical Analysis

Statistical analyses, including generation of graphs and plots, were performed using R versions 3.4.4 and 3.6.0. Significance levels are * = *p* < 0.05, ** = *p* < 0.01, and *** *p* < 0.005 unless otherwise indicated in figure legends. Statistical tests utilized are as indicated in respective text and figure legends.

## SUPPLEMENTAL FIGURE AND TABLE LEGENDS

**Figure S1:**
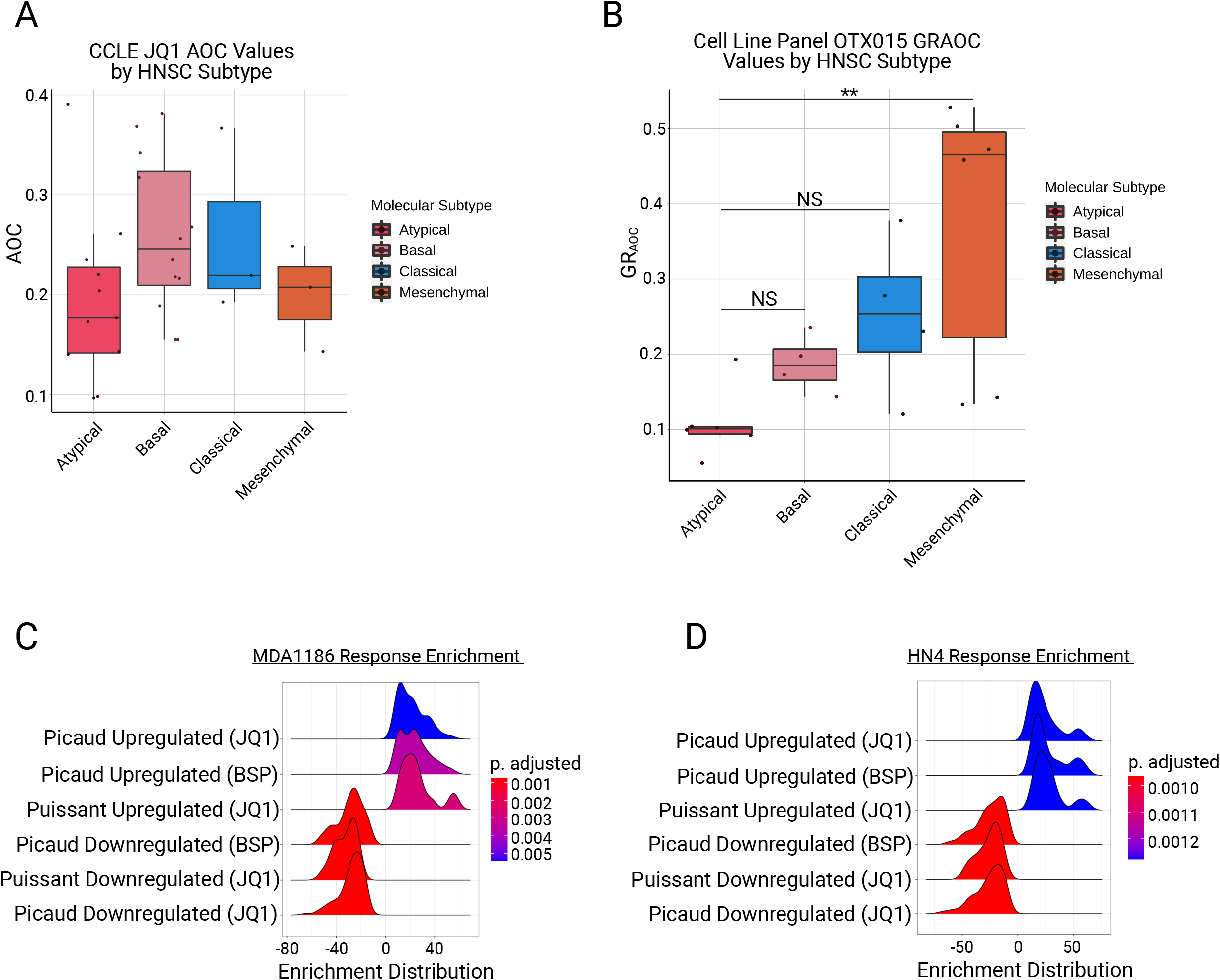
**A**, CCLE-derived JQ1 AOC values grouped by individual HNSC molecular subtype. **B**, Response of HNSC cell lines to the BET inhibitor OTX015, reported as GR_AOC_ to adjust for cell line growth rates and grouped by molecular subtype (** *adj.p* < 0.01). **C, D**, GSEA analysis of **(c)** MDA1186 and **(d)** HN4 response signatures to 24 hours of PLX51107 treatment at their respective GR_50_ values; gene sets for enrichment calculations were generated from previously published BET inhibitor response signatures.

**Table S1:** List of cell lines used for H3K27ac ChIP-seq, their total H3K27ac peak counts, and their total uniquely mapped read counts.

| Cell Line | H3K27ac Peak Count | Uniquely Mapped Read Count |
| --- | --- | --- |
| MDA686TU | 1572 | 21,804,223 |
| MDA1186 | 1645 | 53,936,640 |
| UMSCC4 | 1991 | 18,936,177 |
| TR146 | 2107 | 33,158,077 |
| SCC61 | 2277 | 28,322,650 |
| UMSCC10A | 2530 | 46,547,005 |
| PCI24 | 2738 | 15,519,333 |
| CAL27 | 3759 | 46,177,798 |
| MDA1386LN | 4135 | 35,841,252 |
| MDA1586 | 4689 | 27,684,351 |
| OSC19 | 4840 | 48,540,576 |
| UMSCC19 | 5268 | 23,387,066 |
| 584A2 | 5382 | 36,078,668 |
| 183 | 7649 | 31,378,589 |
| MDA686LN | 7958 | 74,553,208 |
| UMSCC17A | 8527 | 22,288,600 |
| MDA1386TU | 9099 | 15,284,653 |
| PECAPJ34 | 9373 | 18,881,876 |
| UMSCC6 | 10148 | 67,796,124 |
| HN30 | 10284 | 49,741,831 |
| UMSCC25 | 11163 | 24,435,478 |
| MDA886LN | 11297 | 29,432,728 |
| HN5 | 11587 | 50,841,594 |
| HN31 | 12328 | 40,737,765 |
| HN4 | 14825 | 24,428,348 |
| JHU029 | 15037 | 17,917,542 |
| UMSCC14B | 21018 | 35,474,448 |
| MDA1986LN | 22145 | 41,457,423 |

**Table S2:** List of top 20 enriched Hallmark and KEGG pathways and their respective FDRs from genes lists generated by linking AT-unique typical enhancers to their target genes.

Enriched\_pathways\_AT\_unique\_TE
| Gene_set_name | FDR_q_value | Database |
| --- | --- | --- |
| HALLMARK_E2F_TARGETS | 2.47E-09 | Hallmark |
| HALLMARK_MYC_TARGETS_V1 | 7.85E-08 | Hallmark |
| HALLMARK_APOPTOSIS | 7.85E-08 | Hallmark |
| HALLMARK_ADIPOGENESIS | 2.00E-06 | Hallmark |
| HALLMARK_XENOBIOTIC_METABOLISM | 5.41E-06 | Hallmark |
| HALLMARK_TNFA_SIGNALING_VIA_NFKB | 1.46E-05 | Hallmark |
| HALLMARK_P53_PATHWAY | 3.90E-05 | Hallmark |
| HALLMARK_MTORC1_SIGNALING | 1.02E-04 | Hallmark |
| HALLMARK_FATTY_ACID_METABOLISM | 1.67E-04 | Hallmark |
| HALLMARK_APICAL_JUNCTION | 2.11E-04 | Hallmark |
| HALLMARK_G2M_CHECKPOINT | 2.11E-04 | Hallmark |
| HALLMARK_MYC_TARGETS_V2 | 4.11E-04 | Hallmark |
| HALLMARK_HEME_METABOLISM | 4.89E-04 | Hallmark |
| HALLMARK_UV_RESPONSE_UP | 9.55E-04 | Hallmark |
| HALLMARK_HYPOXIA | 9.75E-04 | Hallmark |
| HALLMARK_INTERFERON_GAMMA_RESPONSE | 9.75E-04 | Hallmark |
| HALLMARK_MYOGENESIS | 9.75E-04 | Hallmark |
| HALLMARK_ESTROGEN_RESPONSE_EARLY | 2.17E-03 | Hallmark |
| HALLMARK_OXIDATIVE_PHOSPHORYLATION | 2.17E-03 | Hallmark |
| HALLMARK_DNA_REPAIR | 2.69E-03 | Hallmark |
| KEGG_CELL_CYCLE | 2.03E-05 | KEGG |
| KEGG_SPLICEOSOME | 2.03E-05 | KEGG |
| KEGG_TASTE_TRANSDUCTION | 3.63E-04 | KEGG |
| KEGG_PURINE_METABOLISM | 1.54E-03 | KEGG |
| KEGG_ENDOCYTOSIS | 2.42E-03 | KEGG |
| KEGG_HEMATOPOIETIC_CELL_LINEAGE | 2.42E-03 | KEGG |
| KEGG_FRUCTOSE_AND_MANNOSE_METABOLISM | 4.87E-03 | KEGG |
| KEGG_PEROXISOME | 8.23E-03 | KEGG |
| KEGG_OOCYTE_MEIOSIS | 8.23E-03 | KEGG |
| KEGG_ALZHEIMERS_DISEASE | 9.24E-03 | KEGG |
| KEGG_GLYCOSPHINGOLIPID_BIOSYNTHESIS_GANGLIO_SERIES | 9.24E-03 | KEGG |
| KEGG_MAPK_SIGNALING_PATHWAY | 1.05E-02 | KEGG |
| KEGG_PYRIMIDINE_METABOLISM | 1.26E-02 | KEGG |
| KEGG_RENIN_ANGIOTENSIN_SYSTEM | 1.26E-02 | KEGG |
| KEGG_GLYCOSYLPHOSPHATIDYLINOSITOL_GPI_ANCHOR_BIOSYNTHESIS | 1.26E-02 | KEGG |
| KEGG_NATURAL_KILLER_CELL_MEDIATED_CYTOTOXICITY | 1.26E-02 | KEGG |
| KEGG_RIBOSOME | 1.26E-02 | KEGG |
| KEGG_GLYCEROPHOSPHOLIPID_METABOLISM | 1.28E-02 | KEGG |
| KEGG_PATHWAYS_IN_CANCER | 1.35E-02 | KEGG |
| KEGG_UBIQUITIN_MEDIATED_PROTEOLYSIS | 2.32E-02 | KEGG |

**Table S3:**
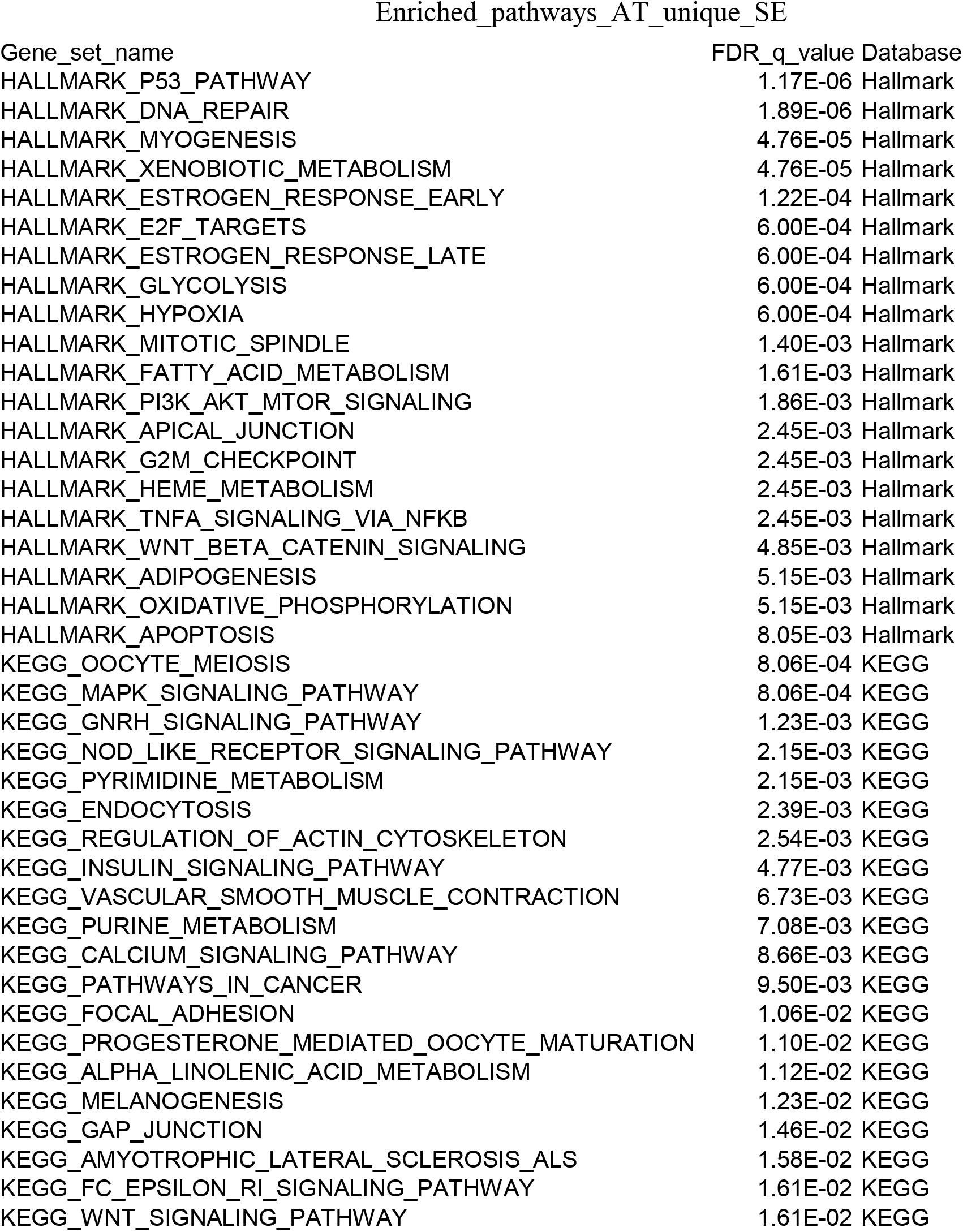
List of top 20 enriched Hallmark and KEGG pathways and their respective FDRs from genes lists generated by linking AT-unique super enhancers (sheet 2) to their target genes.

